# Frequency-dependent Maternal Effects across Species and Environments

**DOI:** 10.1101/225763

**Authors:** Rachel M. Germain, Tess N. Grainger, Natalie T. Jones, Benjamin Gilbert

## Abstract

1. Maternal provisioning of offspring in response to environmental conditions (“maternal environmental effects”) has been argued as ‘the missing link’ in plant life histories. Although empirical evidence suggests that maternal responses to abiotic conditions are common, there is little understanding of the prevalence of maternal provisioning in competitive environments.
2. We tested how competition in two soil moisture environments affects maternal provisioning of offspring seed mass. Specifically, we varied conspecific frequency from 90% (intraspecific competition) to 10% (interspecific competition) for 15 pairs of annual plant species that occur in California.
3. We found that conspecific frequency affected maternal provisioning (seed mass) in 48% of species, and that these responses included both increased (20%) and decreased (24%) seed mass. In contrast, 68% of species responded to competition through changes in per capita fecundity (seed number), which generally decreased as conspecific frequency increased. The direction and magnitude of frequency-dependent seed mass depended on the identity of the competitor, even among species in which fecundity was not affected by competitor identity.
4. *Synthesis*. Our research demonstrates how species responses to different competitive environments manifest through maternal provisioning, and that these responses alter previous estimates of environmental maternal effects and reproductive output; future study is needed to understand their combined effects on population and community dynamics.

## Introduction

The amount of resources available to individual offspring at the propagule stage (seeds or eggs) is maternally controlled, and depends on the mother’s provisioning strategy and resource environment. Changes in propagule size in response to maternal environmental conditions have been shown to have cascading effects on offspring life histories (Segers & Taborsky 2010; Allen 2012) and components of fitness (e.g., germination, dormancy, survival, and reproduction; Westoby *et al*. 1996; Gomez 2004)—a phenomenon known as ‘maternal environmental effects’ and referred to simply as ‘maternal effects’ for brevity henceforth, though other forms exist (Roach & Wulff 1987; Galloway *et al*. 2009). Maternal effects that increase propagule size come at the expense of offspring number (offspring size-number tradeoffs, Charnov & Ernest 2006), but in certain environments, mothers that produce few large offspring can have higher fitness than those that produce many small offspring (Allen, Buckley & Marshall 2008). Because of their clear consequences for offspring fitness in many species, maternal effects have been referred to as ‘the missing link’ between parent and offspring life histories (Donohue 2009), and have remained of interest to evolutionary biologists seeking to understand selection and adaptation for over 50 years (Roach & Wulff 1987; Mousseau & Fox 1998).

The diversity of species-specific maternal effects observed in evolutionary studies Herman & Sultan 2011) implicates the importance of maternal effects for ecological dynamics, such as population persistence and competition. Indeed, the maternal environment can have large effects on trait means and the fitness of whole cohorts of individuals in a population (*i.e*., population growth rates; Galloway & Etterson 2007) that act additively or interactively with offspring environmental conditions (Uller, Nakagawa & English 2013). Because traits and population growth rates mediate how species interact with each other and their environments, maternal effects might act as temporal dimensions to species’ niches that, in addition to responses to current environmental conditions (e.g., Levine & Rees 2004), alter coexistence outcomes; this phenomenon has been demonstrated experimentally in the related field of ‘carryover effects’ of early-life conditions (Van Allen & Rudolf 2015). However, predictions for how maternal effects might influence ecological dynamics cannot be made using existing data from single species experiments because they exclude species interactions, such as competition, that are important to species persistence in multi-species communities (VanAllen & Rudolf 2016).

In competitive environments, maternal effects on offspring provisioning may manifest in response to the relative frequencies of conspecific to heterospecific competitors, even when total density is maintained (Law & Watkinson 1987). In ecological studies, frequency-dependent demographic rates are used to infer the relative strength of competitive interactions within and among species (Levine & HilleRisLambers 2009). Although seeds have been shown to decrease in size in response to increasing plant density (Larios & Venable 2015), tests of maternal effects in response to impacts of different competitive neighborhoods are lacking. Tests that incorporate frequency-dependent effects are particularly relevant, as parallel tests on fecundity are central to understanding species coexistence (Levine & HilleRisLambers 2009) and may be reinforced or counteracted by maternal effects (Germain & Gilbert 2014). Without knowledge of maternal provisioning responses to different competitive neighborhoods, population and community ecologists cannot build intergenerational effects into a broader understanding of population and community dynamics.

Most studies of maternal effects in response to abiotic conditions are conducted in low-competition environments (e.g., Germain & Gilbert 2014) even though organisms rarely occur in the absence of biotic interactors in nature. However, competition might interact with the abiotic environment to affect seed size if competitive interactions alter resource availability or a species’ response to abiotic conditions. For example, competitors may exacerbate maternal effects that are driven by a limiting resource (Stratton 1989), such as soil moisture (Fotelli *et al*. 2001). As a result, current estimates of the prevalence of maternal effects are likely conservative. Species responses to the abiotic environment, conspecific competitors, and heterospecific competitors are necessary components of competition models; predicting the influence of maternal effects on ecological dynamics requires an understanding of how the maternal environment modifies each response.

We test the effects of competition and soil moisture on maternal seed provisioning using 25 annual plant species that occur in the mediterranean-climate regions of the California Floristic Province. The California Floristic Province is characterized by high inter-annual rainfall variability, which determines plant community composition, productivity, and the nature of competitive interactions (Levine, McEachern & Cowan 2011). In variable environments, selection favors plastic responses, such as maternal environmental effects, that offset variability in fitness (Dey, Proulx & Teotónio 2016). The seed stage is important to the life cycle of an annual plant because annual plant populations regenerate entirely each year from the seed bank. Increased seed mass generally provides early growth advantages, allowing individuals to establish prior to the onset of unpredictable hazards, such as drought (Kidson & Westoby 2000), as well as increased competitive ability in productive years (Susko & Cavers 2008).

We competed fifteen pairs of species at six relative frequencies and in two soil moisture environments that simulate wet and dry years. We then quantified the mass and number of seeds produced, and used those data to address three questions: (i) How commonly does maternal provisioning respond to conspecific frequency (i.e., changes in interspecific vs. intraspecific frequency) and how does it compare to seed number responses? (ii) Is maternal provisioning in response to abiotic conditions sensitive to the competitive environment? And (iii) does competitor identity alter the strength and direction of frequency-dependent maternal provisioning? In a previous experiment, we found that ~20% of the same species considered here exhibit maternal effects on seed mass in response to soil moisture conditions in the absence of competition (Germain & Gilbert 2014); we use this earlier study to compare maternal effects in non-competitive and competitive environments.

## Materials and methods

### STUDY SPECIES

To test species responses to competition, we examined maternal effects on seed mass among 15 pairs of annual plant species (25 species total, Table S1) that were competed as part of a previous study (Germain, Weir & Gilbert 2016); species pairs were selected to meet two criteria. First, the 25 species spanned a broad taxonomic range (six angiosperm Orders represented; Table S1), which allowed us to select pairs that represented a range of phylogenetic distances (nine to 170 million years since divergence; phylogeny in Fig. S1). Pairs were selected to represent phylogenetically independent contrasts (i.e., non-overlapping branch lengths), to circumvent phylogenetic pseudoreplication. Second, the selected species overlap in habitat preference (all associate with grassland meadow in mediterranean-type climates) and overlap geographically in California (CalFlora [http://www.calflora.org]); as such, they have the potential to compete in the wild. Additional details about species selection are in Supplementary Methods.

We initially sought a balanced design with 10 species competed twice to test how strongly maternal effects on seed mass were determined by the identity of the competitor, resulting in a total of 20 species pairs and 30 unique species. However, competition was intense enough among 5 species pairs that seeds were not produced and maternal effects could not be quantified, resulting in the design we present here with 15 species pairs and 25 unique species, with five species competed twice. Seeds were obtained from commercial suppliers and an individual donor, and were sourced from natural populations consisting of relatively few generations (most < 3, all < 20; Table S1) prior to our experiments. It is possible that the genetic diversity of our seed populations is low compared to natural populations, though we suspect that this is not the case given the large numbers of individuals used to establish the commercial populations and the small number of generations that have elapsed.

### GREENHOUSE EXPERIMENT

From January to July 2012, we grew the 15 species pairs in competition in a greenhouse under two soil moisture levels (wet vs. dry); see Supplementary Methods for details on growing conditions. Plants in the wet treatment were watered twice as often as those in the dry treatment, with the total water received designed to mimic rainfall in mesic sites during wet and dry years (Germain & Gilbert 2014). The competitive environment was manipulated using a replacement design (Jolliffe 2000), in which seeds of each species pair were sown at six relative frequency ratios (10:60, 20:50, 30:40, 40:30, 50:20 and 60:10 expected germinants) at a common density of 70 individuals. This density is comparable to the seeding density found in annual grasslands (2500 to 5500 plants m^−2^ (Bartolome 1979)). To obtain a density of 70 individuals per pot, we tested each species’ germination rate prior to the experiment and corrected seeding densities based on these rates (i.e. a pot with 60 individuals of one species would receive 60 seeds of a species with 100% germination, or 120 seeds of a species with 50% germination).

For each species pair, we had two replicate pots per combination of soil moisture condition and frequency ratio, for a total of 360 pots of plants that were randomly assigned to a position in the greenhouse. All greenhouse growing conditions were chosen to simulate those typical of annual grassland in mediterranean-climate regions (Germain & Gilbert 2014; Supplementary Methods). We monitored pots daily and collected all seeds produced in each pot as they matured on the parent plants, pooling seed among individuals of the same species in each pot. At the end of the experiment, all seed material produced in each pot was weighed, and a random representative subsample was taken to estimate the average mass per seed and number of seeds produced per plant in a given pot (Supplementary Methods).

Concurrent to this experiment, an additional experiment using the same species was conducted to estimate the impact of soil moisture conditions on maternal provisioning (reported in Germain & Gilbert 2014). Key differences between the previous experiment and the experiment we present here are that each species was grown as singles-species monocultures at low densities (~seven individuals per pot, compared to 70 in the current experiment), meaning that competition was greatly relaxed. We include summary results from Germain and Gilbert (2014) in this paper to compare maternal effects on seed mass to soil moisture conditions in the presence and absence of competition.

### STATISTICAL ANALYSES

Prior to analysis, we transformed the data in two ways to meet model assumptions and facilitate comparisons among species. First, we log transformed seed mass, seed number, and conspecific frequency to minimize heteroscedasticity and linearize seed mass and seed number relationships with conspecific frequency. Second, to allow us to compare species that differ markedly in seed production, we standardized the log-transformed seed mass and seed number data for each species to a mean of zero and unit variance. For simplicity, we henceforth refer to the standardized log-transformed data as ‘seed mass’, ‘seed number’, and ‘frequency’, unless stated otherwise.

We conducted a cross-species analysis to test if species differed in their seed mass and seed number responses to the competitive and soil moisture environments. For these analyses, we used the ‘lmerTest’ R package to run linear mixed effects (LME) models, with either seed mass or seed number as response variables, and species (30 levels), soil moisture (two levels), conspecific frequency (six levels), and their interactions as fixed factors. Because species competing in a pot are not independent of one another, we included ‘Pot ID’ as a random factor in all models to control for the non-independence of species interacting in a single pot and avoid pseudoreplication (Bolker *et al*. 2009). The identity of the competitive pair (e.g., *Lasthenia glabrata* vs. *L. californica*) was also included as a random factor. Following significant species x frequency and species x soil moisture interactions (see Results), we ran species-specific analyses separately to identify species with significant responses to the biotic and abiotic environment. To accomplish this, we used linear models with type II sums of squares to test the effects of conspecific frequency, soil moisture conditions, and their interaction on seed mass and seed number. We used type II sum of squares as opposed to type III sum of squares because the latter prevents interpretation of main effects even in the absence of a significant interaction (Zuur *et al*. 2009). In cases where the interaction was significant, we do not interpret the main effects (e.g., triangle points in Fig. 1). The type of sum of squares used has no effect on the coefficient estimates.

**Fig. 1.**
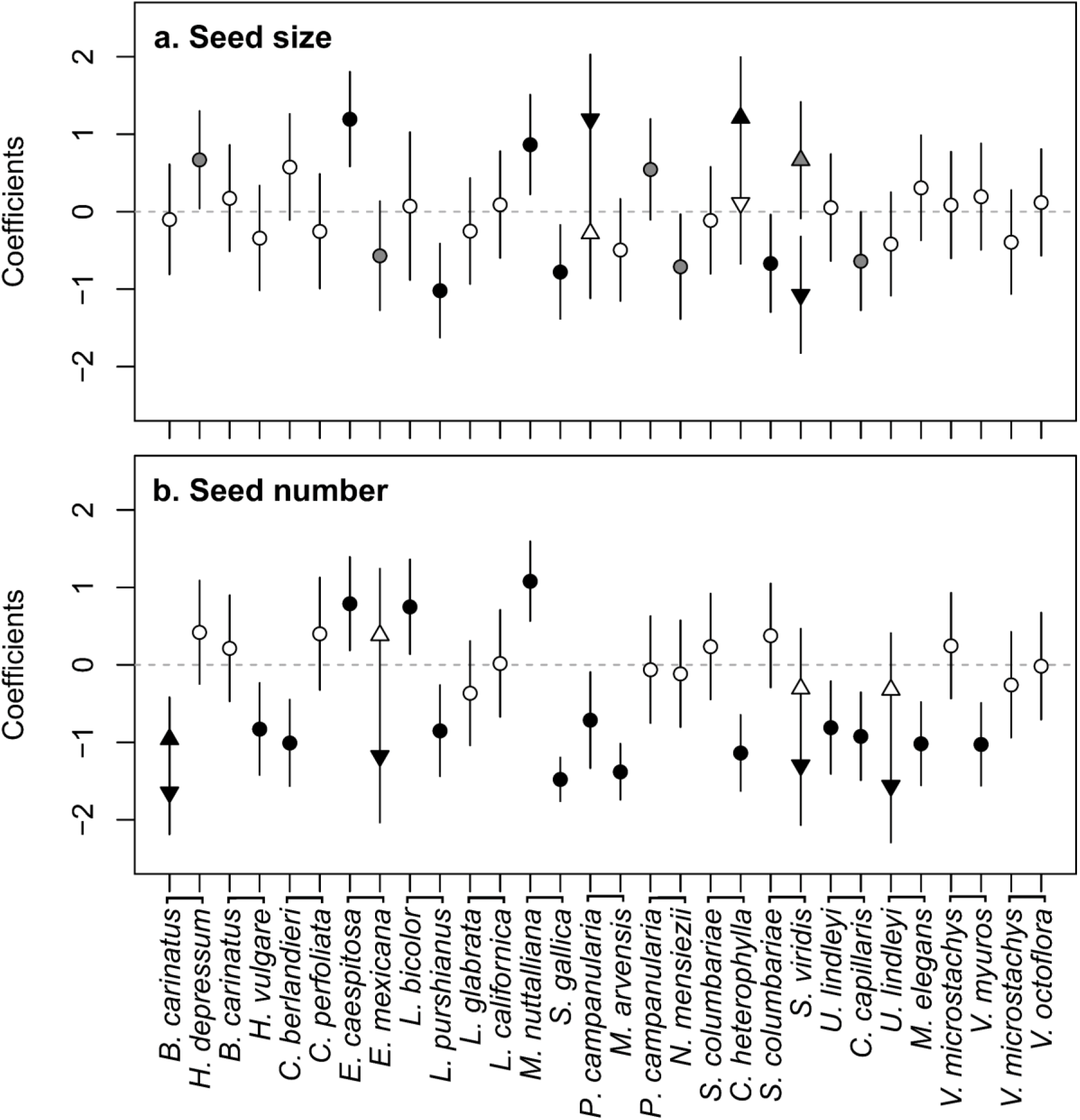
Cross-species comparison of frequency-dependent (a) maternal provisioning of seed mass and (b) seed number responses. Points are slope coefficients (± 2 × standard error) of species responses to the frequency of conspecific competitors, and are shaded black, gray, or white to indicate significant (*P* < 0.05), marginally-significant (*P* < 0.10), or non-significant (*P* > 0.10) slopes, respectively. In most cases, frequency × soil moisture interactions are non-significant, and slopes are averaged across soil moisture environments; when significant, wet (upwards triangle) and dry (downwards triangle) environments are plotted separately. Competitive pairs are delineated by lines connecting species name abbreviations (first letter of genus and species name). See Table S1 for taxonomic and collection information.

We used major axis regression (MAR; R package ‘lmodel2’) to examine the relationships between response variables across species, and tested the significance of these relationships using a Pearson correlation. First, to identify if seed mass and seed number responses are correlated, we performed a MAR with the slopes of species’ seed mass responses and the slopes of species’ seed number responses as variables. Second, we tested whether the presence and absence of competition alters species’ responses to soil moisture conditions. To do this, we first calculated species’ average effect sizes of seed mass responses to contrasting soil conditions (μ_dry_ – μ_wet_) in the presence of competition (i.e. when grown at a density of 70 plants per pot) using Cohen’s d with pooled variance (Hartung, Knapp & Singha 2008); in cases where a species was used in more than one species pair, a single average effect size was used. We then used MAR to examine the relationship between these effect sizes and previously published, identically calculated effect sizes in the absence of competition (i.e. grow at a density of 7 species per pot; Germain & Gilbert 2014). Because these species are small-statured and occur at high densities in the field, we consider the contrast of 70 vs. 7 plants per plot as representative of a competition vs. no competition contrast.

## Results

Of the 25 species included in this study to produce 15 reciprocal competition trials (see Materials and methods), 16 species showed significant (*P* < 0.05) or marginally-significant (*P* < 0.10) maternal effects on seed mass when in competition (Fig. 1a, Fig. S2); we note that 2.5 species are expected to show these results by chance alone given a 10% type 1 error rate. Most species responded to conspecific frequency alone (5 species) or in conjunction with the soil moisture environment in an additive (4 species) or multiplicative (3 species) manner; four species responded to the soil moisture environment but not conspecific frequency. The strength and direction of responses varied among species, as indicated by significant species x frequency (*F_29,464_* = 2.84, *P* < 0.001) and species x soil moisture (*F_29,410_* = 3.19, *P* < 0.001) interactions in our cross-species statistical model (Table S2). Species were similarly likely to increase (5 species) or decrease (6 species) seed mass as conspecific frequency increased (Table S2, Fig. 1a). Overall, our results demonstrate that for this plant community, 64% of species are likely to exhibit maternal effects on seed mass when in competition, and that these responses vary by species with biotic or abiotic conditions.

In contrast to seed mass, 21 of the 25 species showed significant or marginally significant responses through seed number to conspecific frequency (7 species; Fig. 1b), soil moisture conditions (4 species), both additively (6 species), and both interactively (4 species; Table S2). Analogous to seed mass responses, the effect of conspecific frequency on seed number depended on the focal species (significant species x frequency interaction; Table S3, *F_29,716_* = 6.63, *P* < 0.001). However, in contrast to seed mass responses, frequency-dependent seed number was negative for most species (14 out of 17 species; Table S2), and was sensitive to the soil moisture environment (significant frequency x soil moisture interaction; Table S3, *F_1,716_* = 4.49, *P* = 0.034). These seed number responses were positively correlated with the strength and direction of species’ seed mass responses (*r* = 0.28, slope = 0.48, *P* = 0.029; Fig. 2a), even though some species showed opposite seed mass and seed number responses (grey regions of Fig. 2a).

**Fig. 2.**
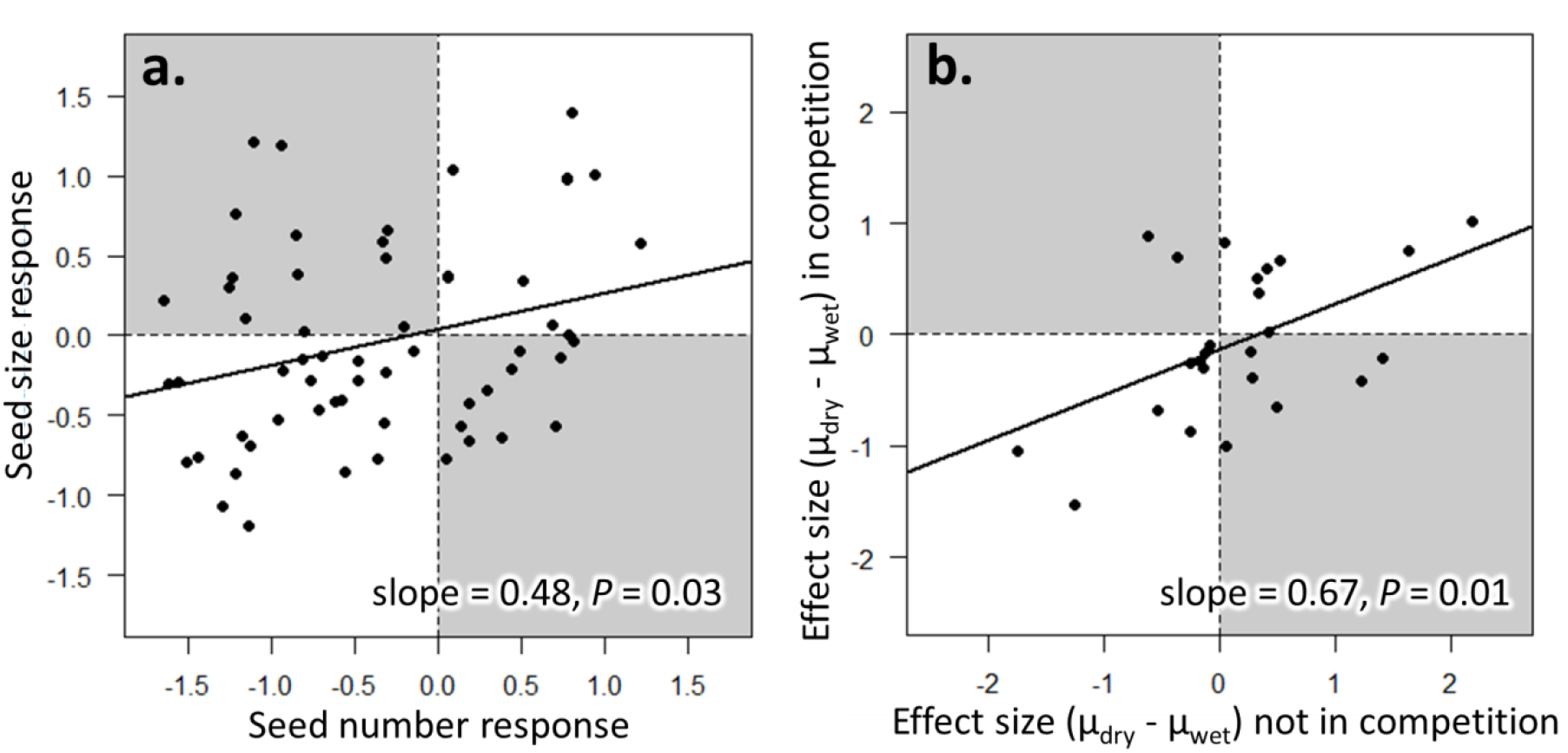
Correlations of (a) seed mass and seed number responses to conspecific frequency (*n* = 60 [15 pairs × two species × two soil moisture environments]), and (b) seed mass responses to soil moisture conditions in the presence and absence of competition (*n* = 25). Each point is a species, and points that fall in the grey zones are species with opposing directions of responses.

The strength and direction of seed mass responses to contrasting soil moisture environments (wet vs. dry) in the presence of competition were positively correlated with those in the absence of competition (*r* = 0.50, *P* = 0.010; Fig. 2b). The slope of this correlation was less than one (major axis regression, slope = 0.67), as many species showed stronger responses to soil moisture when grown in the absence of competition (Fig. 2b). This result was surprising given that, in the current experiment, 44% of species altered seed provisioning in response to soil moisture when in competition, whereas only 21% did so in the absence of competition (Germain & Gilbert 2014). This suggests that the prevalence but not the magnitude of soil moisture-induced maternal effects on seed mass increases in the presence of competition, possibly because the effects of soil moisture are dwarfed by those of conspecific frequency.

## Discussion

There is a substantial body of empirical work investigating maternal effects in response to the abiotic environment (see review by Herman & Sultan 2011), yet responses to biotic interactions remain understudied, particularly in plants (Weiner *et al*. 1997; Larios & Venable 2015). Our results show that changes to the competitive environment can alter maternal provisioning of seed mass and, much like more commonly measured seed number responses (Law & Watkinson 1987), that the strength and direction of maternal provisioning depend on identities of competing species. Below, we discuss how considering interactions between the biotic and abiotic environment allows us to understand how maternal effects are distributed across species in plant communities. Because species interactions frequently altered provisioning strategies, maternal effects likely have important implications for competitive dynamics, species coexistence and diversity; in this vein, we propose new hypotheses for future study.

We detected maternal effects on seed mass in response to the frequency of conspecific competitors in nearly half of the species examined, with negative and positive responses being equally common (Fig. 1a). There are two explanations for the maintenance of species-specific maternal effects that are not mutually exclusive and likely differ in importance among species. First, theory predicts that maternal provisioning strategies should evolve to maximize maternal fitness, either to the benefit or detriment of offspring fitness (Marshall & Uller 2007; Olofsson, Ripa & Jonzen 2009). In that case, the most adaptive strategy will depend on the specific ecologies of focal species (e.g., Sultan, Barton & Wilczek 2009; Germain, Caruso & Maherali 2013). Second, even if the same adaptive strategy is shared by two species, genetic or physiological constraints might result in the evolution of an adaptive or maladaptive strategy in one species but not the other (DeWitt, Sih & Wilson 1998). Though the degree to which species-specific maternal effects reflect evolutionary adaptation or constraint cannot be disentangled in our study, the ecological implications for offspring can be – larger seeds are more competitive and tolerant of environmental stress, whereas smaller seeds are more dispersive and likely to persist longer in the seedbank (Larios & Venable 2015; reviewed in Leishman *et al*. 2000). The life-history tradeoff between competitive ability and dispersal ability, as mediated by seed size, has been hypothesized to play a central role in determining coexistence outcomes (Westoby *et al*. 1996; Jakobsson & Erikson 2000), and might explain the diversity of responses to conspecific frequency observed in our study.

Frequency-dependent seed mass responses were sensitive to the identity of the heterospecific competitor, rather than simply a common response to conspecific frequency. For example, the frequency-dependent seed mass of *Salvia columbariae* was significantly negative when competing with *S. viridis*, but was non-significant in competition with *Collinsia heterophylla*. A simple explanation for this result is that seed mass is sensitive to differences among species’ competitive abilities, yet this mechanism is unlikely given the lack of frequency-dependent seed number responses to either competitor. The presence of seed mass responses in the absence of seed number responses is intriguing, because the latter would typically lead one to conclude that intraspecific and interspecific competition are equivalent (Harpole & Suding 2007). Yet, if competitive interactions were equivalent, we would not expect to see frequency-dependent seed mass responses. Seed mass responses appear to reveal competitive differences among species that are hidden when only seed number responses are considered – as in most ecological studies of plant competition. Previous research in plant monocultures demonstrates that parents produce smaller, more dispersive offspring when neighborhood densities are high (Larios & Venable 2014) – but in multi-species communities, this simple response likely also depends on the relative frequencies of heterospecific competitors.

An open question is whether maternal effects (seed mass responses) act to reinforce or counteract frequency-dependent demographic rates (seed number responses). Although our results suggest that seed mass responses to conspecific frequency generally act to reinforce seed number responses (i.e., they are positively correlated, Fig. 2a), some species clearly show opposite seed mass and seed number responses (points in the grey regions of Fig. 2a). In the context of demographic rates, the direction of frequency dependence can indicate whether competition is more likely to result in coexistence (negative frequency dependence) or exclusion (positive frequency dependence). In our experiment, negative frequency-dependent seed number responses were common among species (fig. 1b), but seed size responses were equally positive and negative (Fig. 1a). Previous research suggests that interspecific and intraspecific variation in seed mass alters several important biological parameters, from dormancy to growth and fecundity (Westoby *et al*. 1996; Eriksson 1999; Germain & Gilbert 2014). Thus, commonly-measured seed number responses (Harpole & Suding 2007) may be insufficient to capture the full impact of competitive interactions in the offspring generation (Ginzburg & Taneyhill 1994; Van Allen & Rudolf 2015). An intriguing avenue for future research are experiments that quantify how much variation in population dynamics is being missed without considering lagged responses to conditions of the maternal generation. An effect of the maternal environment on population demography has been demonstrated previously in response to abiotic conditions (e.g., understory light, Galloway & Etterson 2007), but is not yet understood in competitive environments.

We found important differences in the prevalence of maternal effects in response to soil moisture conditions in competitive and non-competitive environments. Specifically, over twice as many species altered seed provisioning in response to soil moisture in the presence of competition (44% of species; this study, Table S2) compared to in the absence of competition (21% of species; Germain & Gilbert 2014). Almost all tests of maternal effects have been conducted in non-competitive environments, with individuals grown alone (e.g., Aarssen & Burton 1990). Because most plants experience competition in their natural environments, current estimates of the prevalence of maternal effects may be conservative, and most relevant to disturbed environments where plant densities are low. Additionally, competition appears to dampen maternal effects in response to soil moisture (Fig. 2b) despite a threefold increase in the number of species that exhibit such an effect. This indicates that, contrary to our initial expectations, competition does not simply exacerbate maternal provisioning of seed mass in response to soil moisture, but instead appears to alter the nature of soil moisture’s effects on seed mass in some species. This surprising result is likely due to the shift in maternal provisioning that occurs with a change in the identity of the competitor.

An intriguing hypothesis posed by Dyer *et al*. (2010) is that maternal effects might contribute to the invasion success of non-native species. Although our experiment was not specifically designed to test this hypothesis, six of the 25 species used in our trials are not native to California (Table S1), allowing qualitative comparisons. In doing so, we found that frequency-dependent maternal effects are significantly lower among non-native species compared to native species (Fig. 3, Supplementary Methods). Our results suggest that for this system, maternal effects to the competitive environment may be one trait that differentiates native and non-native species and thus could contribute to invasion success. However, we caution that this result is based on limited and unbalanced data (only six non-native species), and could be explored with more species in future studies. For example, we do not have the power to test differences among non-native species that differ in impact, such as naturalized species vs. noxious invaders (Strauss *et al*. 2006; Diez *et al*. 2008), which could explain the considerable overlap in the range of responses among native and non-native species despite significant differences.

**Fig. 3.**
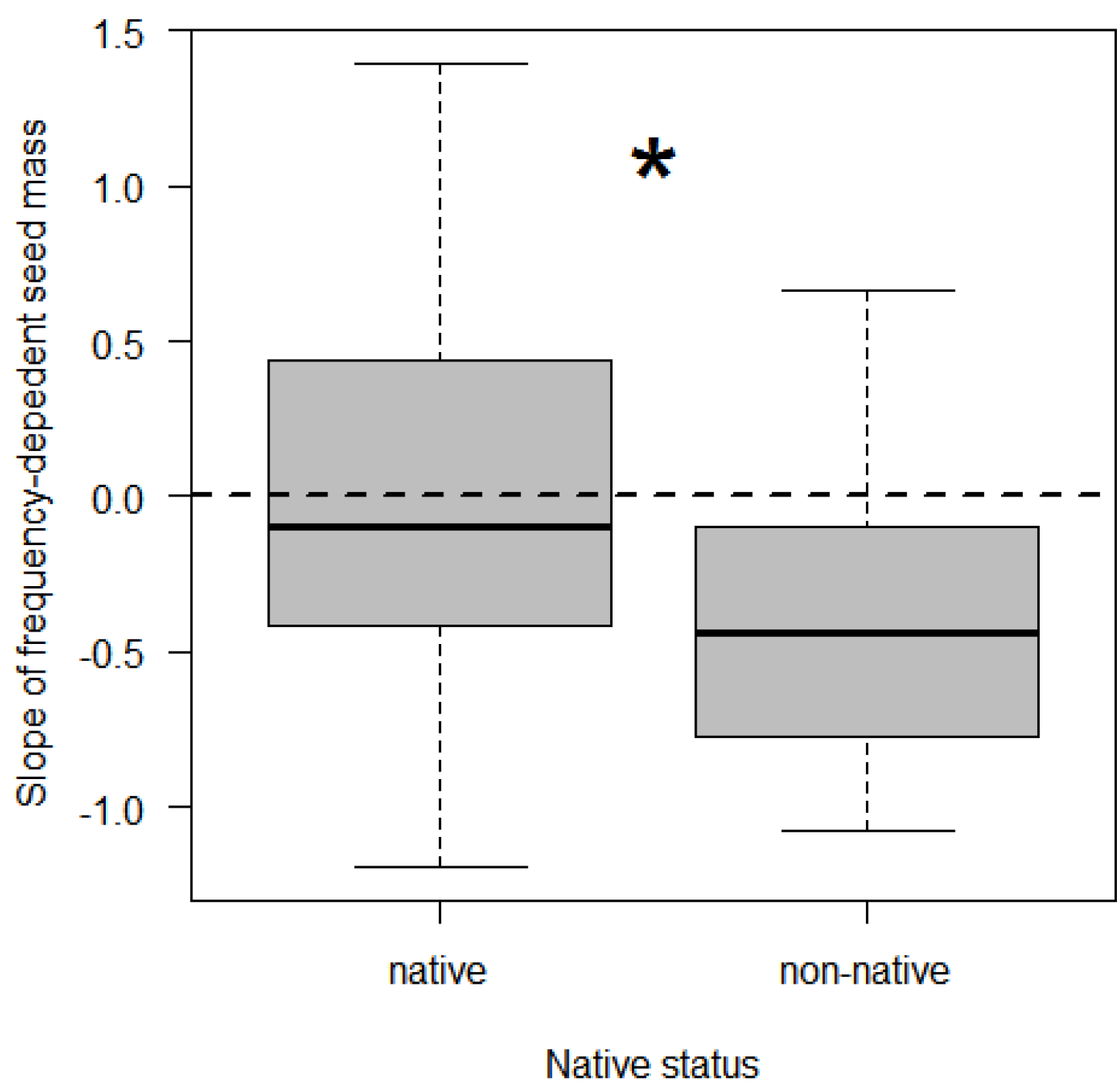
A comparison of frequency-dependent seed mass among native (*n* = 19) and non-native (*n* = 6) species. The dashed horizontal line indicates a slope of 0, and the asterisk indicates a significant difference (*P* = 0.03). See Supplementary Methods for details of the analysis.

Although this study advances our understanding of the importance of competition in structuring maternal provisioning, there are two caveats that should be considered in interpreting our findings. First, we were unable to identify how changes in seed mass translate into differences in offspring performance, due to the logistical infeasibility of the full factorial experiment that would be required to test for longer-term impacts of seed mass. It is possible that maternal effects on seed mass may not persist beyond the seed stage, as some studies have found (e.g., Weiner *et al*. 1997). However, many studies demonstrate their effects on some aspect of post-seed performance, such as germination, dormancy, survival, growth, and fecundity (e.g., Stanton 1984; Gomez 2004; Germain, Caruso & Maherali 2013; see review by Herman & Sultan 2011), especially in competitive environments (Stratton 1989). Second, by focusing on seed mass responses, we likely underestimate the overall prevalence of maternal effects that can manifest in other ways, such as through germination or dormancy rates (Germain & Gilbert 2014), or through epigenetic effects that can alter the offspring phenotype in more complex ways (Herman & Sultan 2011). As such, this study should be viewed as an important first step in characterizing maternal effects in competitive environments that can be used to inform future work.

The study of maternal effects has exciting potential to explain population- and community-level responses to heterogeneous environments (Ginzburg & Taneyhill 1994; Van Allen & Rudolf 2013; Van Allen & Rudolf 2015). Here, we show that current estimates of maternal effects in non-competitive environments are conservative, that competition can alter maternal provisioning of seed mass, and that these maternal effects are fine-tuned to competitive differences among species, which in turn are shaped by the abiotic environment. Our research sheds new light on the complex nature of species interactions, and suggests avenues for future research that would further characterize the full range and impact of maternal effects in ecological communities.

## Acknowledgements

We thank C. Blackford, A. Leale, A. Mushka, B. Hall, and A. Petrie for assistance. Funding was provided by NSERC-CGS (R.M.G. & T.N.G.), OGS (N.T.J.), and NSERC Discovery (B.G.). The authors declare no conflicts of interest.

## Data accessibility

All data will be deposited into the Dryad repository.

